# DNABERT: pre-trained Bidirectional Encoder Representations from Transformers model for DNA-language in genome

**DOI:** 10.1101/2020.09.17.301879

**Authors:** Yanrong Ji, Zhihan Zhou, Han Liu, Ramana V Davuluri

## Abstract

Deciphering the language of non-coding DNA is one of the fundamental problems in genome research. Gene regulatory code is highly complex due to the existence of polysemy and distant semantic relationship, which previous informatics methods often fail to capture especially in data-scarce scenarios. To address this challenge, we developed a novel pre-trained bidirectional encoder representation, named DNABERT, that forms global and transferrable understanding of genomic DNA sequences based on up and downstream nucleotide contexts. We show that the single pre-trained transformers model can simultaneously achieve state-of-the-art performance on many sequence predictions tasks, after easy fine-tuning using small task-specific data. Further, DNABERT enables direct visualization of nucleotide-level importance and semantic relationship within input sequences for better interpretability and accurate identification of conserved sequence motifs and functional genetic variants. Finally, we demonstrate that pre-trained DNABERT with human genome can even be readily applied to other organisms with exceptional performance.

## INTRODUCTION

Deciphering the language of DNA for hidden instructions has been one of the major goals of biological research (1). While the genetic code explaining how DNA is translated into proteins is universal, the regulatory code that determines when and how the genes are expressed varies across different cell-types and organisms (2). Same *cis*-regulatory elements (CREs) often have distinct functions and activities in different biological contexts, while widely spaced multiple CREs may cooperate, resulting in context-dependent use of alternative promoters with varied functional roles (3-6). Such observations suggest existence of polysemy and distant semantic relationship within sequence codes, which are key properties of natural language. Previous linguistics studies confirmed that the DNA, especially the non-coding region, indeed exhibits great similarity to human language, ranging from alphabets and lexicons to grammar and phonetics (7-12). However, how the semantics (i.e. functions) of CREs vary across different contexts (up and downstream nucleotide sequences) remains largely unknown.

In recent years, many computational tools have been developed by successfully applying deep learning techniques on genomic sequence data to study the individual aspects of *cis*-regulatory landscapes, including DNA-protein interactions (13), chromatin accessibility (14), non-coding variants (15) and others. Most methods adopted Convolutional Neural Network (CNN)-based architecture (16). Other tools focus on the sequential characteristic of DNA and attempt to capture the dependency between states by applying Recurrent Neural Network (RNN)-based models, such as Long Short-Term Memory (LSTM) (17) and Gated Recurrent Units (GRU) (18) networks. Several hybrid methods were also proposed to integrate the advantages of the two model architectures (19-21).

To better model DNA as a language, an ideal computational method should (i) globally take all the contextual information into account to distinguish polysemous CREs; (ii) develop generic understanding transferable to various tasks; (iii) generalize well when labeled data is limited. However, both CNN and RNN architectures fail to satisfy these requirements (Figure 1a) (22,23). CNN is usually unable to capture semantic dependency within long-range contexts, as its capability to extract local features is limited by the filter size. RNN models (LSTM, GRU), although able to learn long-term dependency, greatly suffer from vanishing gradient and low-efficiency problem when it sequentially processes all past states and compresses contextual information into a bottleneck with long input sequences. In addition, most existing models require massive amount of labeled data, resulting in limited performance and applicability in data-scarce scenarios, where high quality data with labels is expensive and time-consuming to obtain.

**Figure 1.**
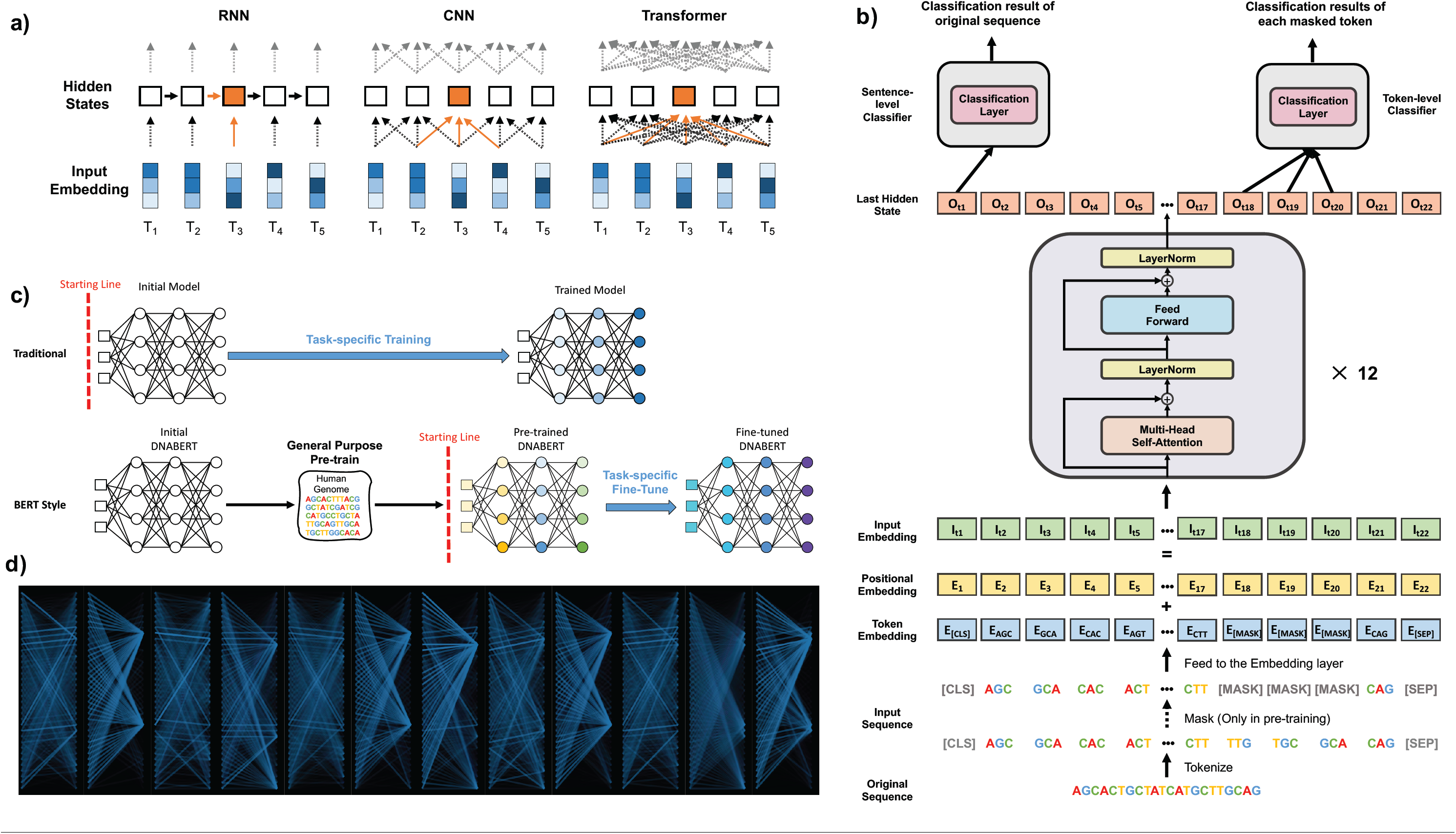
Details of architecture and characteristics of DNABERT model. (a) Differences between RNN, CNN and Transformer in understanding contexts. T1 to 5 denotes embedded tokens which were input into models to develop hidden states (white boxes, orange box is the current token of interest). RNN propagates information through all hidden states, and CNN takes local information in developing each representation. In contrast, Transformers develop global contextual embedding via self-attention. (b) DNABERT uses tokenized k-mer sequences as input, which also contains a CLS token (a tag representing meaning of entire sentence), a SEP token (sentence separator), and MASK tokens (to represent masked k-mers in pre-training). The input passes an embedding layer and is fed to 12 Transformer blocks. The first output among last hidden states will be used for sentence-level classification while outputs for individual masked token used for token-level classification. Et, It and Ot denote the positional, input embedding, and last hidden state at token t respectively. (c) DNABERT adopts general-purpose pre-training which can then be fine-tuned for multiple purposes using various task-specific data. (d) Example of global attention patterns across 12 attention heads showing DNABERT correctly focusing on two important regions corresponding to known binding sites within sequence.

To address the above limitations, we adapted the idea of Bidirectional Encoder Representations from Transformers (BERT) model (24) to DNA setting and developed a first-of-its-kind deep learning method in genomics called DNABERT. DNABERT applies Transformer, an attention-based architecture that has achieved state-of-the-art performance in most natural language processing tasks (25). We demonstrate that DNABERT resolves the above challenges by (i) developing general and transferable understandings of DNA from the purely unlabeled human genome, and utilizing them to generically solve various sequence-related tasks in a “one-model-does-it-all” fashion; (ii) globally capturing contextual information from the entire input sequence with attention mechanism; (iii) achieving great performance in data-scarce scenarios; (iv) uncovering important subregions and potential relationships between different *cis-*elements of a DNA sequence, without any human guidance; (v) successfully working in a cross-organism manner. Since the pre-training of DNABERT model is resource-intensive (about 25 days on 8 NVIDIA 2080Ti GPUs), as a major contribution of this study, we provide the source code and pretrained model on GitHub for future academic research.

## MATERIAL AND METHODS

### The DNABERT model

DNABERT takes a set of sequences represented as k-mer tokens as input (Figure 1b). Each sequence is represented as a matrix *M* by embedding each token into a numerical vector. Formally, DNABERT captures contextual information by performing the multi-head self-attention mechanism on *M*:

where

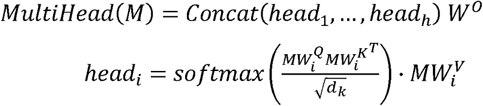

*w*° and 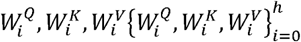 are learned parameters for linear projection. *head* calculates the next hidden states of *M* by first computing the attentions scores between every two tokens and then utilizing them as weights to sum up lines in 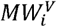. *MultileHead()* concatenates results of *h* independent *head* with different set of 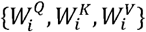. The entire procedure is performed *L* times with *L* being number of layers. More details are presented in Methods section.

DNABERT adopts *pre-training—fine-tuning* scheme (Figure 1c). In the general-purpose pre-training step, DNABERT learns basic syntax and semantics of DNA via self-supervision. For each 510-length sequence in human genome, we randomly mask regions of *k* contiguous tokens that constitute 15% of the sequence and let DNABERT to predict the masked sequences based on the remainder, which ensures ample training examples. We pre-trained DNABERT with cross-entropy loss: 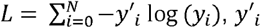 and *y*_′*i*_ being the ground-truth and predicted probability for each of *N* classes. The pre-trained DNABERT model can be fine-tuned with task-specific training data for applications in various sequence- and token-level prediction tasks. We fine-tuned DNABERT model on three specific applications – prediction of promoters, transcription factor binding sites (TFBSs) and splice sites – and benchmarked the trained models with the current state-of-the-art tools.

### Training of DNABERT model

#### Tokenization

Instead of regarding each base as a single token, we tokenized a DNA sequence with the k-mer representation, an approach that has been widely used in analyzing DNA sequences. The k-mer representation incorporates richer contextual information for each deoxynucleotide base by concatenating it with its following ones. The concatenation of them is called a k-mer. For example, a DNA sequence “ATGGCT” can be tokenized to a sequence of four 3-mers: {ATG, TGG, GGC, GCT} or to a sequence of two 5-mers: {ATGGC, TGGCT}. Since different k leads to different tokenization of a DNA sequence. In our experiments, we respectively set k as 3,4,5 and 6 and train 4 different models: DNABERT-3, DNABERT-4, DNABERT-5, DNABERT-6. For DNABERT-k, the vocabulary of it consists of all the permutations of the k-mer as well as 5 special tokens: [CLS] stands for classification token; [PAD] stands for padding token, [UNK] stands for unknown token, [SEP] stands for separation token and [MASK] stands for masked token. Thus, there are 4^*k*^+5 tokens in the vocabulary of DNABERT-k.

#### Pre-training

Following previous works (24,26,27), DNABERT takes a sequence with a max length of 512 as input. As illustrated in Figure 1b, for a DNA sequence, we tokenized it into a sequence of k-mers and added a special token [CLS] at the beginning of it (which represents the whole sequence) as well as a special token [SEP] at the end (which denotes the end of sequence). In the pre-training step, we masked contiguous k-length spans of certain k-mers (to prevent overfitting, total ∼15% of input sequence), while in the fine-tuning, we skipped the masking step and directly fed the tokenized sequence to the Embedding layer. We generated training data from human genome via 2 approaches: direct non-overlap splitting and random sampling, with length of the sequence between 5 and 510. We pre-trained DNABERT for 120k steps with a batch size of 2000. In this first 100k steps, we masked 15 percent of k-mers in each sequence. In the last 20k steps, we increased the masking rate to 20 percent. The learning rate was linearly increased (a.k.a, warm-up) from 0 to 4e^-4^ in the first 10k steps and then linearly decreased to 0 after 200k steps. We stopped the training procedure after 120k steps since we found the loss curve show a sign of plateauing. We used the same model architecture as the BERT base, which consists of 12 Transformer layers with 768 hidden units and 12 attention heads in each layer, and the same parameter setting across all the four DNABERT models during pre-training. We trained each DNABERT model with mixed precision floating point arithmetic on machines with 8 Nvidia 2080Ti GPUs. More details included in Supplementary Methods.

#### Fine-tuning

For each downstream application, we started from the pre-trained parameters and fine-tuned DNABERT with task-specific data. We utilized the same training tricks across all the applications, where the learning rate was first linear warmed-up to the peak value and then linear decayed to near 0. We utilized AdamW with fix weight decay as optimizer and employed dropout to the output layer. We splitted training data into training set and developing set for hyperparameter tuning. For DNABERT with different k, we slightly adjusted the peak learning rate. The detailed hyperparameter settings were listed in Table S5.

### Visualizing attention on sequence via DNABERT-viz

With the help of self-attention mechanism, DNABERT is naturally suitable for locating and deciphering upstream or downstream regulatory regions in genome. To directly visualize the important regions on input sequence that the model uses as evidence to make the final classification decision, we developed a new method (DNABERT-viz) for direct visualization of nucleotide-level scores. Specifically, the self-attention mechanism naturally serves as a scoring approach for individual component of input sequence. Formally, let *q\** be the query vector of the “CLS” token, which is a special symbol appended in front of each sequence and is used for final classification, and let *d* be the dimension of *q\**. Let *k*_*j*_ be the key vector for the *j*-th k-mer token, *j* ∈{1,… *J*}, where is the number of tokens in input sequence. Then, the attention score of each embedded k-mer token over all the attention heads *H* is the sum of softmax

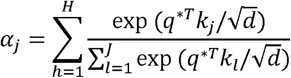

Essentially, we are extracting the attention of the “entire sequence” on the k-mer subsequences and use it as an importance measure. To convert the attention score from k-mer to individual nucleotide level, for a particular nucleotide, the scores for all k-mers that contain it were averaged. Attention for individual nucleotide was then plotted as heatmap for direct visualization. For visualization of token-level self-attention over attention heads (e.g. “context plot” in Figure 4e), we applied the attention-head view in BertViz tool (28) with the display of attention restricted to that greater than a user-specified cutoff.

**Figure 2.**
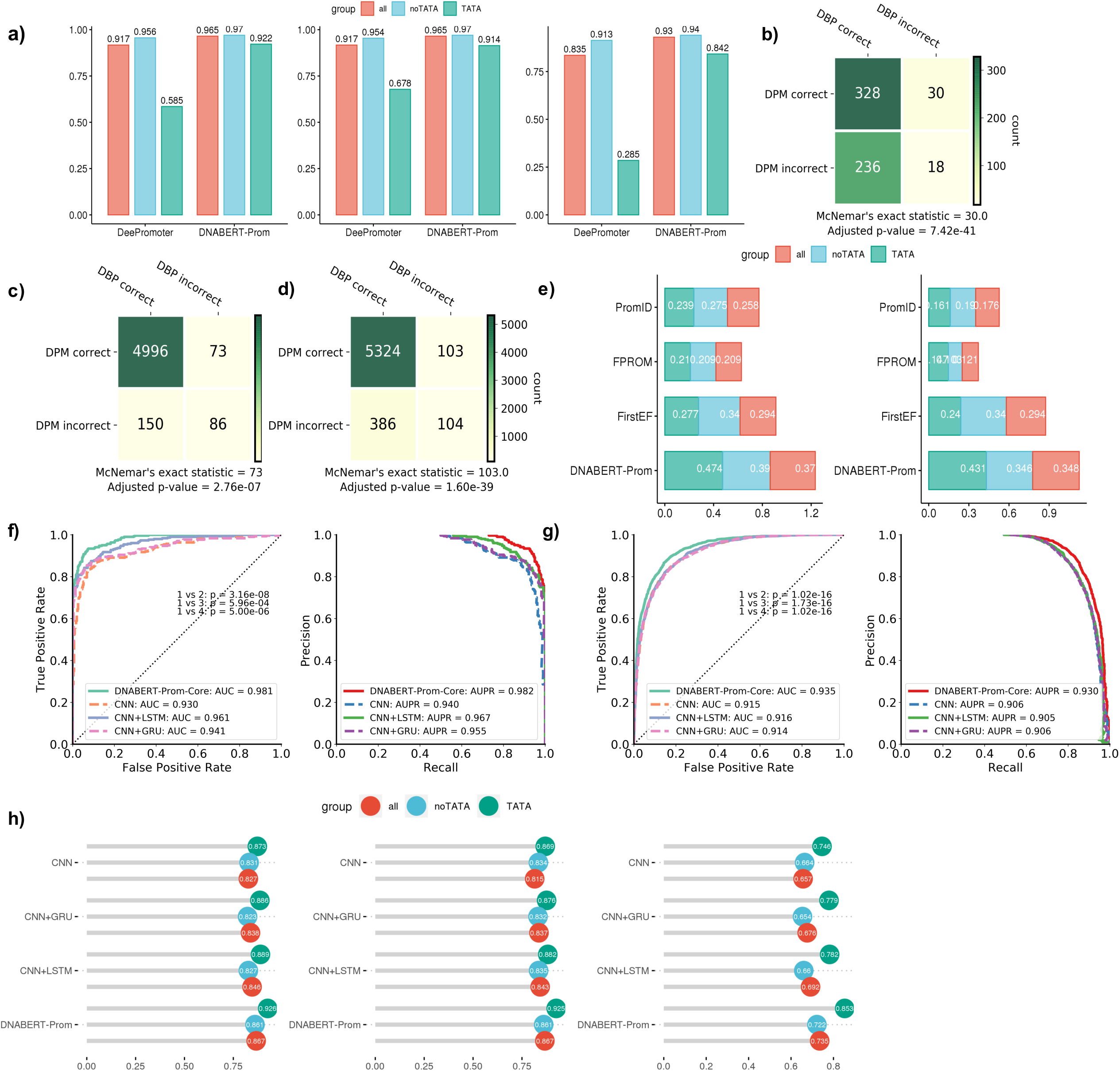
DNABERT significantly outperforms other models in identifying promoter regions. (a) (Left to right) accuracy, F1 and MCC of prom-300 prediction in TATA, noTATA and combined datasets. (b-d) McNemar’s exact test between DNABERT-Prom (DBP) and DeePromoter (DPM) in TATA, noTATA and combined datasets respectively. (e) Stacked barplot showing F1 (left) and MCC (right) of Prom-scan predictions in different settings. (f-g) ROC (left) and precision-recall (PR) curves (right) on TATA (f) and noTATA (g) datasets with adjusted p-values from Delong test. (h) (Left to right) accuracy, F1 and MCC of core promoters prediction in TATA, noTATA and combined datasets.

**Figure 3.**
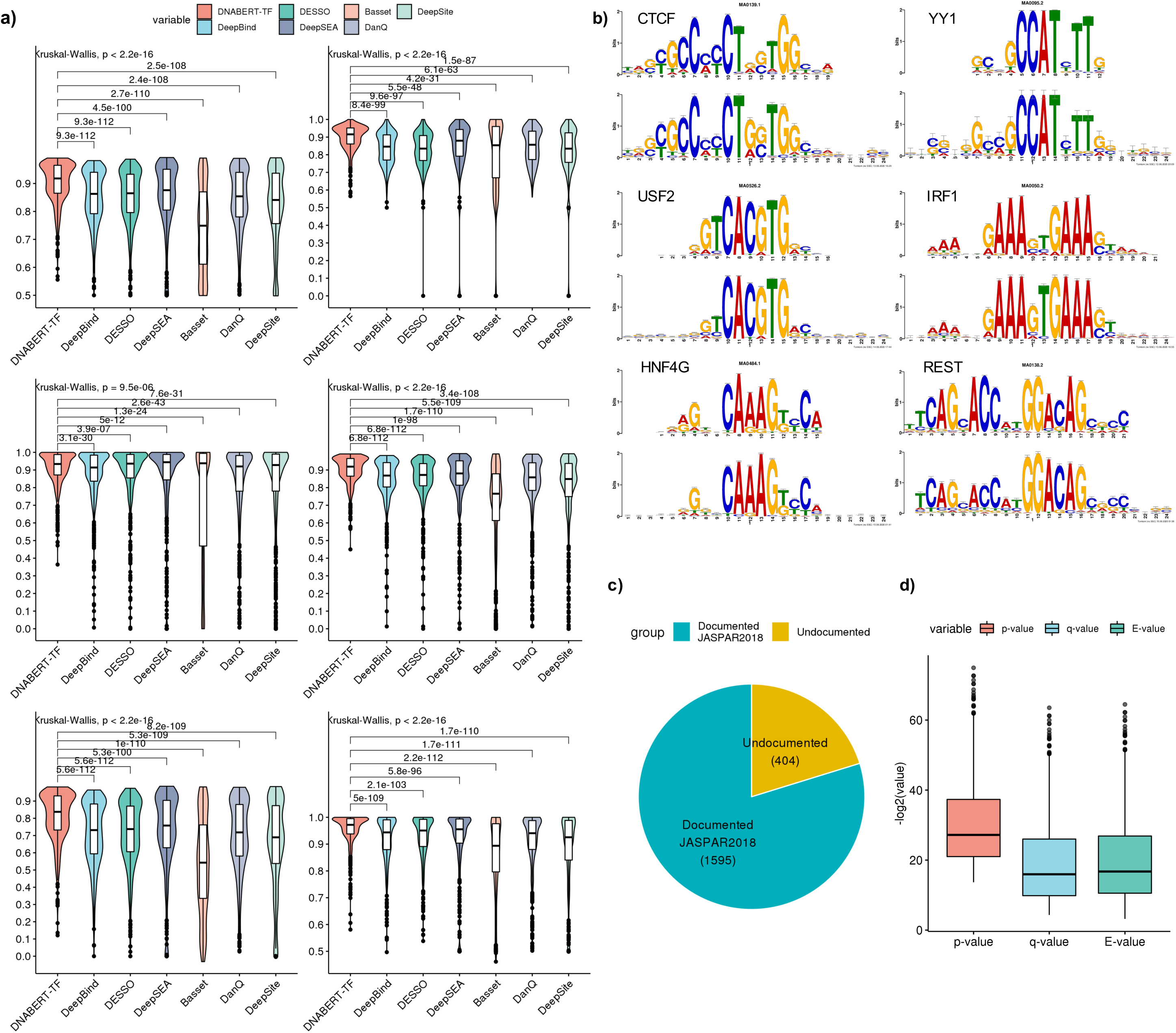
DNABERT accurately identifies TFBSs. (a) Violin plots showing ccuracy (top left), precision (top right), recall (middle left), F1 (middle right), MCC (bottom left) and AUC (bottom right) of TFBS prediction with ENCODE 690 ChIP-Seq datasets. Pairwise comparison using Wilcoxon one-sided signed-rank test (n=690) and adjusted p-values using Benjamini-Hochberg procedure were shown. Global hypothesis testing across all models done by Kruskal-Wallis test (n=690). (b) Selected motifs found by DNABERT and validated in JASPAR2018 database. (Top) TOMTOM documented motifs; (bottom) DNABERT predicted motifs. (c) Summary statistic of documented vs. undocumented motifs in JASPAR2018 database as identified by DNABERT model. (d) –log2(p-value), -log2(q-value) and – log2(E-value) from TOMTOM motif comparison analysis.

**Figure 4.**
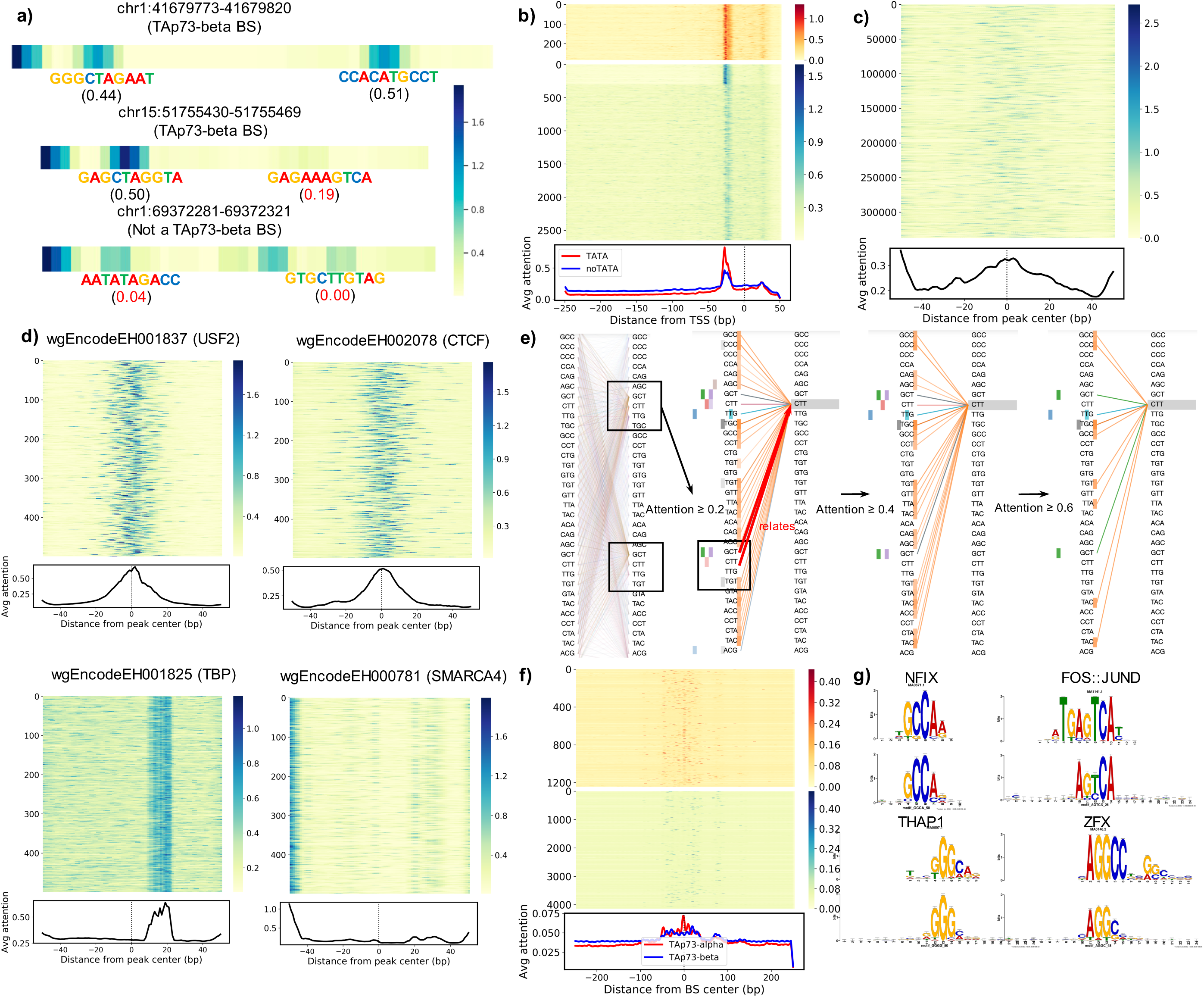
Visualizations of attention and context by DNABERT-viz. (a) Attention maps of 2 example ChIP-Seq-validated TAp73-beta binding sites (top, middle) and one non-binding site (bottom). Numbers below represent binding scores previously predicted by P53Scan. (b, c, f) Attention landscapes of (b) TATA (top) and noTATA (bottom) promoters in Prom-300 test set and (c) all center 101bp of ChIP-Seq peaks from ENCODE 690 dataset. (d) Example attention landscapes for individual ENCODE 690 dataset. USF2, CTCF and TBP are of good quality while SMARCA4 has concerned quality. (e) Attention-head (context) plots of a p53 binding site. (left) sentence-level self-attention across all heads; (middle left, middle right, right) attention of the “CTT” token within one of the important regions, with only attention ≥ 0.2, 0.4 and 0.6 shown respectively. Heatmap on the left shows the corresponding attention head. (g) Selected short motifs identified as enriched in TAp73-alpha as compared to TAp73-beta.

### Motif analysis with DNABERT

In order to extract biologically important motifs enriched in a set of sequences, we developed a motif analysis tool accompanying DNABERT-viz module. Specifically, we first identified contiguous high attention regions within input sequences based on user-defined cutoff conditions. In our analysis, only the regions with (1) attention > mean of attention within the sequence; (2) attention > 10 times minimum of attention within the sequence; and (3) has a minimum length of 4 will be selected. These attention regions were used as preliminary motif instances. Next, we assumed that the random variable *x* representing number of positive sequences containing a motif instance follows a Hypergeometric distribution *x* ∼ *Hypergeom*(*N,K,n*) (29):

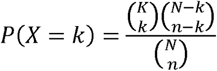

Where *N* stands for total number of sequences, *K* stands for number of positive sequences, *n* stands

for number of sequences containing the specific motif instance and *k* stands for number of positive sequences with the motif instance. A hypergeometric test against *H*_0_ the motif instance is overrepresented/enriched in positive sequences could then be constructed. We applied Aho-Corasick algorithm for efficient multi-pattern matching and computation of hit counts *n* and *k* for a particular motif and we restricted our algorithm to count only once if multiple hits were found in one sequence. We performed hypergeometric test and applied Benjamini-Hochberg procedure for multiple testing correction and filtered motif instances with adjusted p-value < 0.005. Since the model is not guaranteed to place high attention on entire region of a motif instance, some of the significant motif instances identified in fact belong to same motifs. Thus, we merged the similar motif instances by performing pairwise alignment between all pairs. To keep the integrity of all motif instances, we specifically prohibited our aligner from introducing internal gaps. We declared the success of an alignment if the score exceeded the maximum between (required length for contiguous region – 1) and (half of minimum length of the pair). In case of any tie, we merged the motif instance twice with the corresponding two best aligned motifs. In order to convert into a position-weight matrix (PWM) type format, all sequences within a motif were required to be of same length. Therefore, we extracted fixed-length window (24 in our analysis) around center of each motif instance we identified. Finally, we removed motifs with less than 3 instances. The final motif files were converted into Weblogo format and compared with JASPAR2018 validated motifs using TOMTOM command-line version (30).

### Identifying effects of genetic variants using DNABERT

In order to quantify the effects of genetic variants on prediction *p*(*s*) of a sequence (*s*_1_,…,*s*_n_), we substituted the locus of interest *s*_*i*_ with base *b*_*j* ∈_ {*A,T,C,G*},*b*_*j*_ ≠ *S*_*i*_ and recomputed the prediction *p*(*s′*). The genetic effect of the mutation *b*_*j*_ at locus *i* can therefore be calculated using the predicted probabilities. We computed the score change as the differences between the probabilities based on the suggestions in (13): *S* = Δ*p* = (*p*(*s*′) - *p*(*s*) max (*p*(*s*′),*p*(*s*)), where the max term was added to amplify the strong effects of certain genetic variants; as well as the log odds ratio log_2_*OR* = 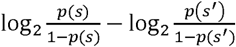 as used in (15), which reflects the association between events “being classified as a positive” and “having the particular genetic variant”. A larger log odds ratio (>0) indicates that the event of being classified as positive is more likely to occur in the reference group i.e. no variant; and *vice versa*. A log odds ratio = 0 indicates no association between the two events. In the case of splice sites prediction, where the output is probability of three classes, the score for donor and acceptor was calculated separately against the non-splice site case. To find variants with functional importance, we downloaded dbSNP release 153 (both GRCh37/hg19 for ENCODE 690 and GRCh38/hg38 for other analysis) containing approximately 700 million short genetic variants and mapped with identified high attention regions within a set of sequences (31). The alleles at corresponding locations were altered and the mutated sequences subjected to predictions. dbSNP Common variants with large absolute change score or logOR score, derived from the predictions on original and mutated sequences and defined as score greater than the average of scores for all variants, were queried in ClinVar (32), GRASP (33), and NHGRI-EBI GWAS Catalog (34), which contain both clinical and functional (GWAS and eQTL, etc.) variants.

### Data access and preprocessing

All datasets used in this study were publicly available and collected from different sources. For pre-training of DNABERT, we downloaded the reference human genome GRCh38.p13 primary assembly from GENCODE Release 33 (35), removed all sequences gaps and/or unannotated regions (sequence regions with “N”) and extracted 5 to 510-nt-long sequences as training data (details in the “pre-training” section below). For promoter prediction, we obtained human TATA and non-TATA promoter data from Eukaryotic Promoter Database (EPDnew) (36) using the provided API EPD selection tool (https://epd.epfl.ch/human/human_database.php?db=human). We extracted −249∼+50 bp sequences around TSS for the Prom-300 setting, −34∼+35 bp for Prom-core setting, and 1001-bp-long scans for Prom-scan setting. For TFBS prediction, we retrieved the ENCODE 690 ChIP-seq profiles from UCSC genome browser that covers 161 TFs in 91 human cell lines (37,38) (http://hgdownload.cse.ucsc.edu/goldenPath/hg19/encodeDCC/wgEncodeAwgTfbsUniform/) and extracted the center 101 bp as TFBS-containing sequences following other benchmarking studies. For analysis of p53, TAp73-alpha and TAp73-beta binding sites, we obtained respective ChIP-seq peaks from Gene Expression Omnibus (GEO) GSE15780 (39) and used those as target regions merged with the p73/p53 binding sites predicted by our P53Scan program (40). For mouse TFBS we downloaded mouse ENCODE ChIP-seq data from Stanford/Yale (41) stored on UCSC genome browser (http://hgdownload.soe.ucsc.edu/goldenPath/mm9/encodeDCC/wgEncodeSydhTfbs/). Finally, for splice sites analysis, we extracted 400-bp-long sequences around the donor and acceptor sites again using GRCh38.p13 genome. More detailed data preprocessing steps for each individual task were covered in Supplementary Methods.

## RESULTS

### DNABERT-Prom effectively predicts proximal and core promoter regions

Predicting gene promoters is one of the most challenging bioinformatics problems. We began by evaluating our pre-trained model on identifying proximal promoter regions. To fairly compare with existing tools with different sequence length settings, we fine-tuned two models, named DNABERT-Prom-300 and DNABERT-Prom-scan, using human TATA and non-TATA promoters of 10,000 bp length, from Eukaryotic Promoter Database (EPDnew) (36). We compared DNABERT-Prom-300 with DeePromoter (42) using −249 to 50 bp sequences around TSS as positive examples, randomly selected 300 bp-long, TATA-containing sequences as TATA negative examples, and dinucleotide-shuffled sequences as non-TATA negative examples (Supplementary Methods). We compared DNABERT-Prom-scan with currently accessible methods, including recent state-of-the-art methods PromID (43), FPROM (44), and our previous software FirstEF (45), using sliding window-based scans from 10,000 bp-long sequences. To appropriately benchmark with PromID under same setting, we used 1,001 bp-long scans, which exceed the length capacity of traditional BERT model. Hence, we developed DNABERT-XL specifically for this task (Supplementary Methods). We used same evaluation criteria used in PromID by scanning sequences and overlapping predictions with −500 to +500 bp of known TSS. The resulting 1,001 bp sequences with ≥ 50% overlap to −500 to +500 bp of TSS were deemed as positives and the remaining as negatives. For PromID and FPROM, the test set was directly input for evaluation. In contrast, FirstEF first generates genome-wide predictions, which were then aligned to the positive sequences.

DNABERT-Prom outperformed all other models by significantly improved accuracy metrics regardless of different settings (Figure 2). Specifically, for prom-300 setting TATA promoters, DNABERT-Prom-300 exceeded DeePromoter in accuracy and MCC metrics by 0.335 and 0.554, respectively (Figure 2a). Similarly, we observed significantly improved performance of DNABERT-Prom in both non-TATA and combined cases (Figure 2b-d). Meanwhile, the prom-scan setting is intrinsically more difficult as the classes are highly imbalanced, so all the tested baseline models performed poorly. Among the baselines, FirstEF achieved the best performance with an F1-score of 0.277 for TATA, 0.377 for non-TATA and 0.331 for combined datasets (Figure 2e). However, DNABERT-Prom-scan achieved F1-score and MCC that largely surpassed FirstEF. Next, we evaluated our model’s predictive performance on core promoters, a more challenging problem due to reduced size of the sequence context. We used 70 bp, centered around TSS, of the Prom-300 data and compared with CNN, CNN+LSTM and CNN+GRU. DNABERT-Prom-core clearly outperformed all the three baselines across different datasets (Figure 2f-h), clearly demonstrating that DNABERT can be reliably fine-tuned to accurately predict both the long proximal promoters and shorter core promoters, relying only on nearby sequence patterns around the TSS region.

### DNABERT-TF accurately identifies transcription factor binding sites

NextGen sequencing (NGS) technologies have facilitated genome-wide identification of gene regulatory regions in an unprecedented way and unveiled the complexity of gene regulation. An important step in the analyses of *in vivo* genome-wide binding interaction data is prediction of TFBS in the target *cis-*regulatory regions and curation of the resulting TF binding profiles. We thus fine-tuned DNABERT-TF model to predict TFBSs in the ChIP-seq enriched regions, using 690 TF ChIP-seq uniform peak profiles from ENCODE database (37) and compared with well-known and previous published TFBS prediction tools, including DeepBind (13), DeepSEA (15), Basset (14), DeepSite (46), DanQ (20) and DESSO (47). DNABERT-TF is the only method with both mean and median accuracy and F1-score above 0.9 (Figure 3a, 0.918 and 0.919), greatly exceeding the second best competitor (DeepSEA, Wilcoxon one-sided signed-rank test, n=690, adjusted p = 4.5×10^−100^ and 1×10^−98^ for mean). Other tools made many false positive (FP) and false negative (FN) predictions in certain experiments, resulting in even less satisfactory performance, when comparing the mean due to skewed distribution (Table S1). Several tools achieved comparable performance with DNABERT in finding the true negatives (TN) for experiments using high-quality data, yet performed poorly when predicting on low-quality experimental data. In contrast, even on low-quality data, DNABERT achieved significantly higher recall than other tools (Figure 3a, middle left). Meanwhile, DNABERT-TF made much fewer FP predictions than any other model regardless of the quality of the experiment (Figure 3a, top right). These conclusions are further supported by benchmarking using only subset of ChIP-seq profiles with limited number of peaks, where DNABERT-TF consistently outperformed other methods (Figure S1).

To evaluate whether our method can effectively distinguish polysemous *cis*-regulatory elements, we focused on p53 family proteins (which recognize same motifs) and investigated contextual differences in binding specificities between TAp73-alpha and TAp73-beta isoforms. We overlapped p53, TAp73-alpha and TAp73-beta ChIP-seq peaks from Gene Expression Omnibus (GEO) dataset GSE15780 with binding sites predicted by our P53Scan program (39,40) and used the resulting ChIP-seq-characterized BS (∼35 bp) to fine-tune our model. DNABERT-TF achieved near perfect performances (∼0.99) on binary classification of individual TFs (Table S2). Using input sequences with a much wider context (500 bp), DNABERT-TF effectively distinguished the two TAp73 isoforms with an accuracy of 0.828 (Table S2). In summary, DNABERT-TF can accurately identify even very similar TFBSs based on the distinct context windows.

### DNABERT-viz allows visualization of important regions, contexts and sequence motifs

To overcome the common “black-box” problem, deep learning models need to maintain interpretability, while exceling in performance in comparison to traditional methods. Therefore, to summarize and understand important sequence features on which fine-tuned DNABERT models base classification decisions on, we developed DNABERT-viz module for direct visualization of important regions contributing to the model decision. We demonstrate that DNABERT is naturally suitable for finding important patterns in DNA sequences and understanding their relationship within contexts due to the attention mechanism, thus ensuring model interpretability.

Figure 4a shows the learned attention maps of three TAp73-beta response elements, where DNABERT-viz accurately determines both positions and scores of TFBS predicted by P53Scan in an unsupervised manner. We then aggregated all heatmaps to produce attention landscapes on test sets of Prom-300 and ENCODE 690 TF. For TATA promoters, DNABERT consistently put high attention upon −20 to −30 bp region upstream of TSS where TATA box is located, while for majority of non-TATA promoters a more scattered attention pattern is observed (Figure 4b). Such pattern is also seen in TF-690 datasets, where each peak displays a distinct set of high attention regions, most of which scattered around the center of the peaks (Figure 4c). We specifically looked into examples of individual ChIP-seq experiments to better understand the attention patterns. Most high-quality experiments show enrichment of attention either around the center of the ChIP-seq peaks or on TFBS region (Figure 4d). In contrast, low-quality ones tend to have dispersed attention without strongly observable pattern, except the high attention only at the beginning of sequences, which is likely due to model bias (Figure 4d, bottom right).

We next extended DNABERT-viz to allow for direct visualization of contextual relationship within any input sequence (Figure 4e). For example, the leftmost plot shows global self-attention pattern of an input sequence in the p53 dataset, where individual attentions from most k-mer tokens over all heads correctly converge at the two centers of the dimeric BS. We can further infer the interdependent relationship between the BS with other regions of input sequence by observing which tokens specifically paid high attention to the site (Figure 4e, right). Among attention heads, the orange one clearly discovered hidden semantic relationship within context, as it broadly highlights various short regions contributing to attention of this important token CTT. Moreover, three heads (green, purple and pink) successfully relate this token with the downstream half of the dimeric binding site, demonstrating contextual understanding of the input sequence.

To extract conserved motif patterns across many input sequences, we applied DNABERT-viz to find contiguous high-attention regions and filtered them by hypergeometric test (Methods). The resulting significant motif instances were then aligned and merged to produce position-weight matrices (PWMs). By applying TOMTOM program (30) on the discovered motifs from ENCODE 690 dataset and compared with JASPAR 2018 database, we found that 1,595 out of 1,999 motifs discovered successfully aligned to validated motifs (Figure 3c, q-value < 0.01). Motifs identified are overall of very high quality illustrated by strong similarity to the documented motifs (Figure 3b&d).

We finally applied DNABERT-viz to understand important factors in distinguishing binding sites of TAp73-alpha from beta isoforms. The attention landscape indeed shows many short regions differentially enriched between two isoforms, with alpha having higher attention concentrated at center and beta more scattered into the contexts (Figure 4f). Many strong motif patterns extracted were not aligned to JASPAR database except for a few highlighting unknown relationship (Figure 4g). Importantly, differential crosstalk between c-Fos, c-Jun and TAp73-alpha/beta isoforms contributes to apoptosis balance (48), and DNABERT-viz successfully captured this relationship. To conclude, DNABERT can attain comparable interpretability as CNN-based models in a more straightforward way while greatly surpassing them in prediction performance.

### DNABERT-Splice accurately recognizes canonical and non-canonical splice sites

Predicting splice sites is essential for revealing gene structure and understanding alternative splicing mechanisms. Nevertheless, the presence of both GT-AG-containing non-splice site sequences, and non-canonical splice sites without the dinucleotides, creates difficulty for accurate identification (49). Recently, SpliceFinder (49) successfully addressed this issue by reconstructing a dataset via recursive inclusion of previously misclassified false positive sequences. To compare with SpliceFinder performance on identical benchmark data, we iteratively rebuilt the same dataset with donor, acceptor and non-splice site classes. We also performed comparative analysis with multiple baseline models. As expected, all models performed well on initial dataset as the task is oversimplified, although DNABERT-Splice still achieved the best (Table S3). We, then, compared DNABERT-Splice with all baselines using a reconstructed dataset that includes “adversarial examples” (Figure 5a). This time, the predictive performance of the baseline models greatly dropped, while DNABERT-Splice still achieved best accuracy of 0.923, F1 of 0.919 and MCC of 0.871, with AUROC and AUPRC significantly greater than other models (Figure 5b and c), which was also supported by Mcnemar’s exact test (Figure S2&3). Furthermore, DNABERT-Splice again outperformed all models when predicting on an independent test set containing held-out sliding-window scans from our iterative training process (Table S4). We also examined the attention landscape to elucidate on how model made classification decision (Figure 5d). Surprisingly, DNABERT-Splice showed globally consistent high attention upon intronic regions (downstream of donors and upstream of acceptors), highlighting the presence and functional importance of various intronic splicing enhancers (ISEs) and silencers (ISSs) acting as CREs for splicing (50).

**Figure 5.**
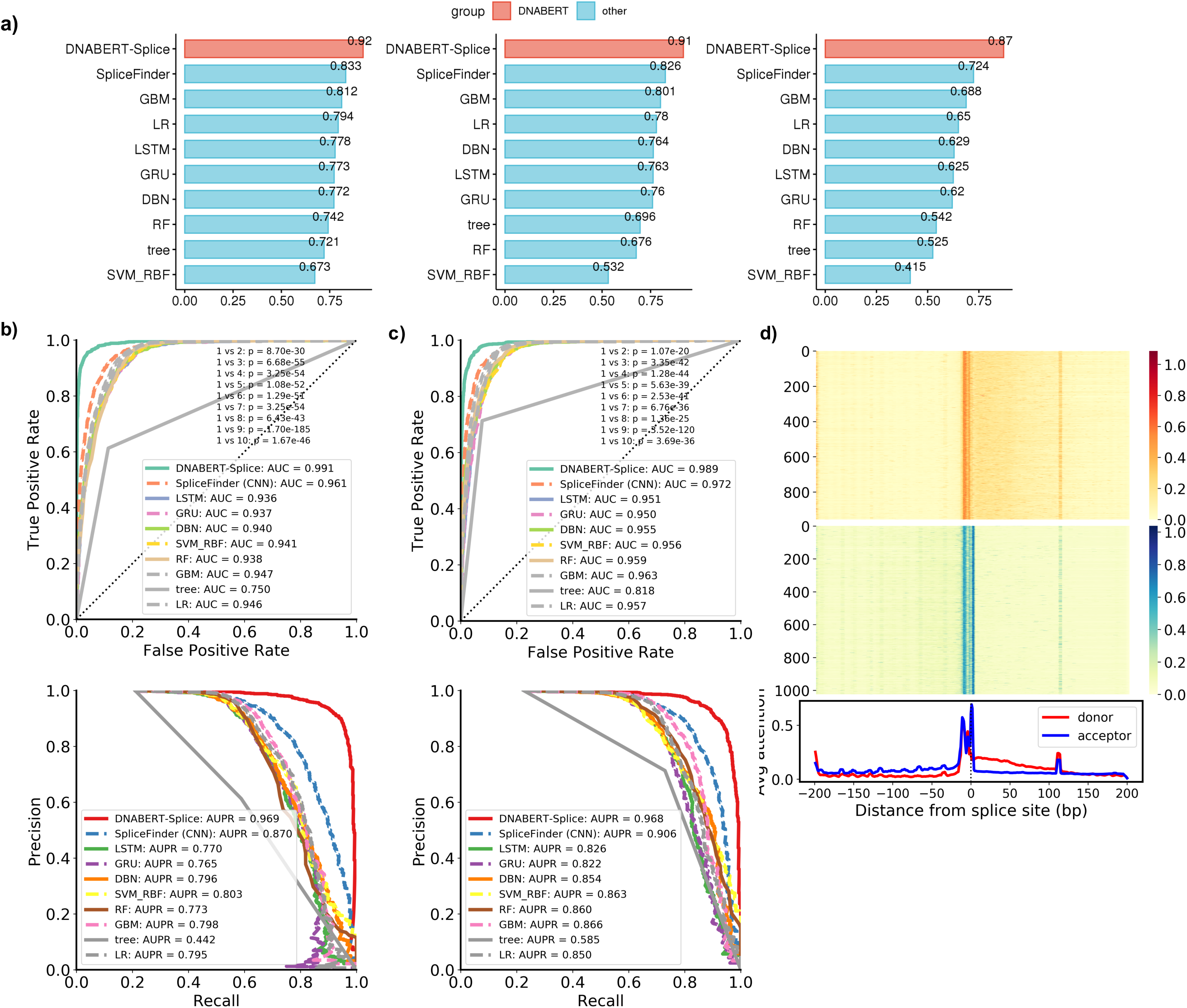
DNABERT significantly outperforms other models in finding splice sites. (a) (Left to right) multiclass accuracy, F1 and MCC of splice donor and acceptor prediction. GBM: gradient boosting; LR: logistic regression; DBN: deep belief network; RF: random forest; tree: decision tree; SVM_RBF: support vector machine with radial basis function kernel. (b, c) ROC (top) and PR curves (bottom) on splice donor (b) and acceptor (c) datasets with adjusted p-values from Delong test. (d) Attention landscape of splice donor (top) and acceptor (bottom).

### Identifying functional genetic variants with DNABERT

We applied DNABERT to identify functional variants using around 700 million short variants in dbSNP (31). Specifically, we selected only those variants that are located inside DNABERT predicted high-attention regions and repeated the predictions, using sequences with altered alleles (Methods). Candidate variants resulting in significant changes in prediction probability were queried in ClinVar (32), GRASP (33), and NHGRI-EBI GWAS Catalog (34). In Prom-300 dataset, we found 24.7% and 31.4% of dbSNP Common variants we identified using TATA and non-TATA promoters are present in at least one of the three databases (Table S6). We present some example functional variants that we found using ENCODE 690 ChIP-seq datasets (Figure 6a-c). Figure 6a shows a rare, pathogenic 4bp deletion completely disrupts a CTCF BS within *MYO7A* gene in ECC-1 cell line. This deletion is known to cause Usher Syndrome, an autosomal recessive disorder characterized by deafness, although the relationship with CTCF is to be determined (51). Similarly, Figure 6b depicts how a rare single nucleotide variant (SNV) at initiator codon of *SUMF1* gene, which leads to multiple sulfatase deficiency (52), simultaneously disrupts a YY1 BS with unknown functional consequences. In Figure 6c, a common risk variant of pancreatic cancer at intronic region of *XPC* gene also greatly weakens CTCF BS (53). In all examples, DNABERT consistently shows highest attention at/around the variants of interest.

**Figure 6.**
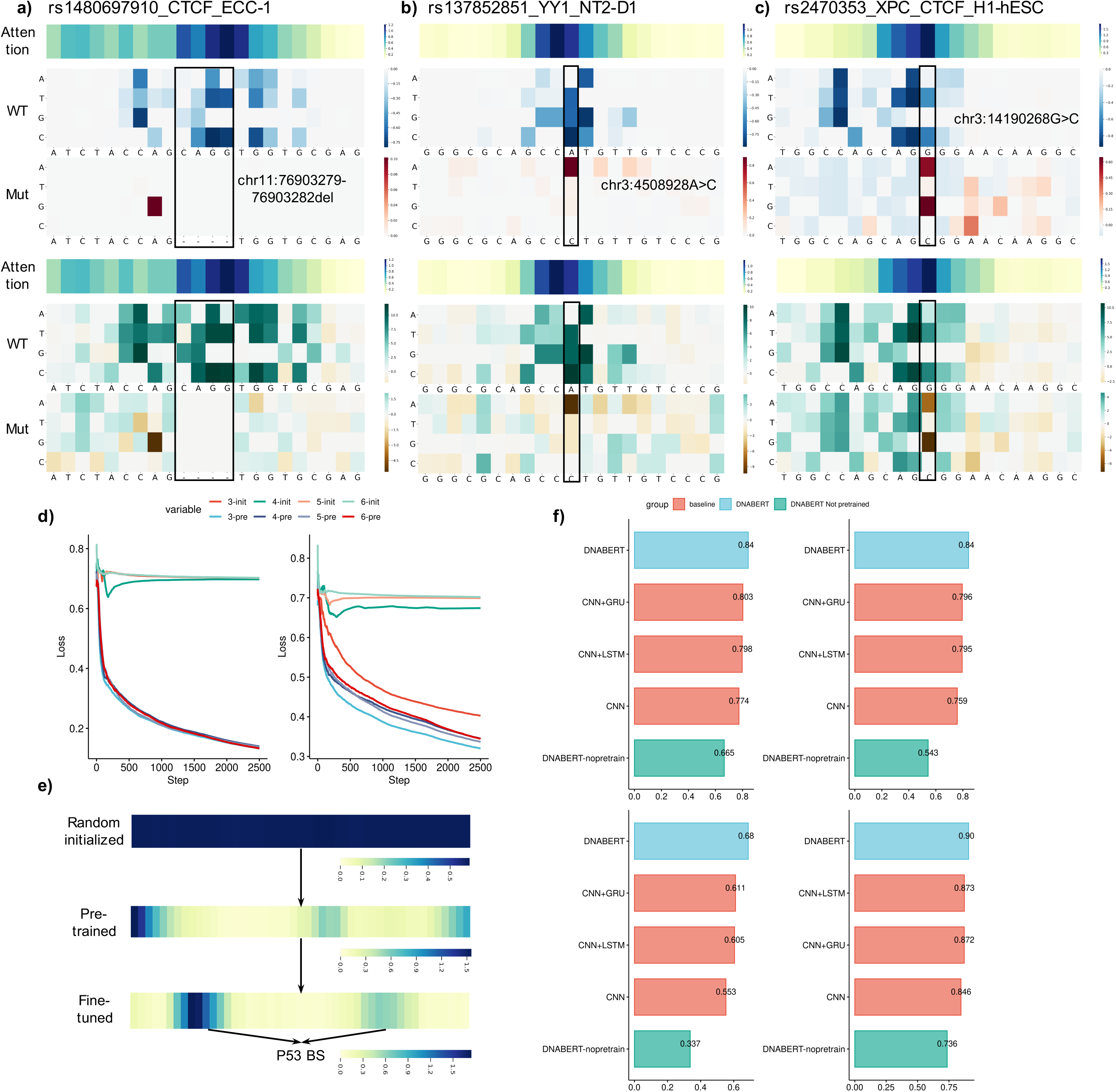
DNABERT identifies functional genetic variants, and pretraining is essential and can be generalized. (a-c) Mutation maps of difference scores (top 3) and log-odds ratio scores (logOR, bottom 3). Each mutation map contains the attention score indicating importance of the region (top), scores for wild-type (WT, middle) and scores for mutant (mut, bottom). (Left to right) a rare deletion within a CTCF binding site inside MYO7A gene in ECC-1 cell line completely disrupts the binding site; a rare single-nucleotide variant (SNV) at initiator codon of SUMF1 gene also disrupts YY1 binding site (5‘-CCGCCATNTT-3’); a common intronic SNP within XPC gene weakens CTCF binding site and is associated with pancreatic cancer. (d) Fine-tuning loss of pre-trained (pre) vs random initialized (init) DNABERT on Prom-300 (left) and Prom-core (right). (e) p53 attention map for random initialized (top), pre-trained (middle) and fine-tuned (bottom) DNABERT model. (f) Mean Accuracy (top left), F1 (top right), MCC (bottom left) and AUC (bottom right) across 78 mouse ENCODE datasets.

### Pre-training substantially enhances performance and generalizes to other organisms

Lastly, we investigated the importance of pre-training based on performance enhancement and generalizability. When comparing training loss of pre-trained DNABERT-prom-300 with randomly initialized ones under same hyperparameters, pre-trained DNABERT converges to a markedly lower loss, suggesting that randomly initialized models get stuck at local minima very quickly without pre-training, as it ensures preliminary understanding of DNA logic by capturing distant contextual information (Figure 6d). Similarly, randomly initialized DNABERT-prom-core models either remain completely untrainable or exhibit suboptimal performance. An examination of attention maps reveals the gradual comprehension of input sequence (Figure 6e). Since separate pre-training of DNABERT for different organisms can be both very time-consuming and resource-intensive, we also evaluated whether DNABERT pre-trained on human genome can be also applied on other mammalian organisms. Specifically, we fine-tuned DNABERT pre-trained with human genome on 78 mouse ENCODE ChIP-seq datasets (41) and compared with CNN, CNN+LSTM, CNN+GRU and randomly initialized DNABERT. Pre-trained DNABERT significantly outperformed all baseline models (Figure 6f), showing the robustness and applicability of DNABERT even across a different genome. It is well known that although the protein-coding regions between human and mouse genomes are ∼85% orthologous, the non-coding regions only show ∼ 50% global similarity (54). Since TFBS mostly locate within the non-coding region, DNABERT model successfully transferred learned information from one genome to a much less similar genome with very high tolerance to the dissimilarities. This demonstrates that the model correctly captured common deep semantics within DNA sequences across organisms. The evaluations above demonstrates the essentiality of pre-training and guarantees extensibility of the pre-trained model for efficient application in numerous biological tasks across different organisms.

## DISCUSSION

In this study, we demonstrated that DNABERT, being a first-of-its-kind BERT model in DNA sequence analysis, achieved state-of-the-art performance across various downstream tasks by largely surpassing existing tools. Using an innovative global contextual embedding of input sequences, DNABERT tackles the problem of sequence specificity prediction with a “top-down” approach by first developing general understanding of DNA language via self-supervised pretraining and then applying it to specific tasks, in contrast to the traditional “bottom-up” approach using task-specific data. These characteristics of DNABERT ensures that it can more effectively learn from DNA context with great flexibility adapting to multiple situations, and enhanced performance with limited data. In particular, we also observed great generalizability of pre-trained DNABERT across organisms, which ensures the wide applicability of our method without the need for separating pre-training.

A major contribution of this work is to release the pre-trained model, and we expect DNABERT to also apply to other sequence analyses tasks, for example, determining *cis*-regulatory elements from ATAC-seq (55) and DAP-seq (56). Further, since RNA sequences differs from DNA sequences only by one base (thymine to uracil), while the syntax and semantics remain largely the same, our proposed method is also anticipated to perfectly apply to Cross-linking and immunoprecipitation (CLIP-seq) data for prediction of binding preferences of RNA-binding proteins (RBPs) (57).

Although direct machine translation on DNA is not yet possible, the successful development of DNABERT shed light on this possibility. As a successful language model, DNABERT correctly captures the hidden syntax, grammar and semantics within DNA sequences and should perform equally well on *Seq2seq* translation tasks once token-level labels become available. Meanwhile, other aspects of resemblance between DNA and human language beyond text (e.g. alternative splicing and punctuation) highlights the need to combine data of different level for more proper deciphering of DNA language. To summarize, we anticipate that DNABERT can bring new advancements and insights to the bioinformatics community by bringing advanced language modeling perspective to gene regulation analyses.

## DATA AVAILABILITY

The source code, pretrained and finetuned model for DNABERT are available at GitHub (https://github.com/jerryji1993/DNABERT).

## ACKNOWLEDGEMENT

We thank Dr. Manoj Kandpal for his help with the initial processing of GEO datasets and reading of the manuscript.

