## Supplementary Materials for "DNABERT: pre-trained Bidirectional Encoder Representations from Transformers model for DNA-language in genome"

**Supplementary Methods**

**Transformer**

Transformer-based models have achieved state-of-the-art performance in various important tasks such as machine translation and question answering. A Transformer model consists of an encoder and a decoder and has an entirely attention-based architecture [1]. Since the BERT model is essentially a multi-layer Transformer encoder, we specifically focus on the encoder part which consists of two sub-layers: the multi-head self-attention layer and the fully connected feed-forward layer. A skip/residual connection structure and layer-wise normalization are incorporated around each sub-layer to greatly facilitate training. The key innovation within encoder is the multi-head self-attention layer which allows the model to associate all the relevant words in a context to encode a specific word better and develop the “contextual understanding” in different aspects.

The attention function computes an output based on a query (*q*) and a set of key-value pairs (*K, V*). The dot-product attention calculates attention scores by multiplying query with each key and using the products as weights to sum all the values, which can be calculated as:

$$Attention\left( q,K,V \right)= softmax\left( qK^{T} \right)\cdot V=\sum_{i} \frac{exp(qK_{i})}{\sum_{j} exp(qK_{j})}\cdot V_{i}$$

Where $K_{i}$ and $V_{i}$ stands for the *i-th* key-value pair. Intuitively, the dot-product between $q$ and $K_{i}$ measures the relevance of $V_{i}$ in representing *q* (how much the model should attend to). Moreover, if we pack a set of queries into a matrix *Q*, and divide the dot-product by a scaling parameter $\sqrt{d_{k}}$, the scaled dot-product attention is calculated as follows:

$$Attention\left( Q,K,V \right)=softmax\left( \frac{QK^{T}}{\sqrt{d_{k}}} \right)\cdot V$$

Here, $d_{k}$ is the dimension of the keys. The self-attention is a special case of scaled dot-product attention, where *Q, K* and *V* come from the same place. Practically, instead of setting Q=K=V and calculate attention scores as $Attention\left( Q,K,V \right)$, it is beneficial to linearly project *Q, K* and *V* with different and learnable parameters $W^{Q}, W^{K}$ and $W^{V}$. Then, attention scores can be calculated as $Attention\left( QW^{Q},KW^{K},VW^{V} \right)$. By performing this independently for multiple times, concatenating all the attention scores and once again projected, we get multi-head attention as:

$$MultiHead\left( Q,K,V \right)=Concat\left( {head}_{1}, \ldots,{head}_{h} \right) W^{O}$$

where	${head}_{i}=$ $Attention\left( QW_{i}^{Q},KW_{i}^{K},VW_{i}^{V} \right)$

where $W_{i}^{Q}$, $W_{i}^{K},$ $W_{i}^{V}$ and $W^{O}$ are learnable parameters. Thus, the input and output of each Transformer layer are both matrix of the same shape. Each line of the matrix stands for the representation of its corresponding token.

Transformer differs from Recurrent Neural Network (RNN) and Convolutional Neural Network (CNN) in calculating each token's hidden states. For RNN, the recurrent unit captures global contexts by taking input from both the previous layer and the previous time step. However, even with the LSTM architecture, RNN still suffers from gradient vanishing when the input sequences are extra-long. Also, its recurrent nature prevents it from parallel computing. The convolutional neural network is parallelizable, but it can only capture the local context. Instead of using the convolutional and recurrent mechanism, Transformer performs the self-attention mechanism on all the representations from the previous layer to calculate the hidden states. On the one hand, it efficiently captures the global contexts and effectively overcome the gradient vanishing issue. One the other hand, it is straightforward to parallelly compute the multi-head attention on a large scale. Thus, the Transformer is regarded as an excellent candidate for large-scale, long sequence modeling that effectively addresses the aforementioned problems of CNNs and RNNs.

The raw input of the Transformer is a sequence of tokens. To feed linguistic tokens into a model, the first thing is to transform each of them into a numerical representation. This representation is always called Token Embedding. To achieve this, a vocabulary and an Embedding layer are essential. The vocabulary is a set that contains all the possible linguistic tokens, and the embedding layer $E$ is essentially a learned $n \mathrm{by} m$ matrix where $n$ equals to the vocabulary size and $m$ equals to the customized embedding size. Each line of the matrix stands for the token embedding of a unique token in the vocabulary. The same tokens will be assigned the token embedding. The Embedding layer directly maps a linguistic token in the vocabulary to an $m$-dimensional vector. Besides, since the self-attention mechanism does not take sequence order into account, to ensure that the model recognizes the order of the input tokens, a position-dependent vector called positional embedding is assigned to each token. The sum of the token embedding and positional embedding is the final input of the Transformer.

**Bidirectional Encoder Representation Transformer (BERT)**

The advent of the BERT model leads the natural language processing research to a new era by introducing a paradigm of pre-training and fine-tuning. Models are first pre-trained on a massive amount of unlabeled data to learn the general rules and relationships and then fine-tuned on task-specific data to learn to perform specific tasks.

As mentioned above, the BERT model is essentially an Embedding layer follows by multiple Transformer Encoder layers. Unlike the vanilla Transformer model, the input sequence of BERT could be either a single sentence or the concatenation of two sentences. The sentence here stands for an arbitrary span of continuous text. This setting enables the BERT model to handle either a single sentence of a pair of sentences as input, which enriches its versatility in dealing with down-stream tasks. A special classification token ([CLS]) is always added to the first position of the input, whose last hidden state is considered as the aggregated representation of the sequence. A special token ([SEP]) separates two sentences in an input sequence.

In the pre-training stage, the input is always the concatenation of two sentences. For each input sequence, 15% tokens are randomly masked, and the model is trained to predict the masked parts based on the rest. This task is usually called masked language model. In addition, the model is asked to predict whether the second sentence is the actual next sentence of the first sentence or not. The losses of these two tasks are summed up to optimize the model.

To decrease the mismatch between pre-training and fine-tuning, for each masked token, (i) with 80% probability; it will be replaced by a special token ([MASK]), (ii) with 10% probability; it will be replaced by a random token, (iii) with the other 10% probability; it will keep the same. The pre-training is performed in a self-supervised fashion. There is no need for any human labeling, yet the training procedure is supervised. This nature allows developers to painlessly involve a massive amount of data for training. In the fine-tuning stage, the model is initialized with pre-trained parameters, and further trained with task-specific data. Fine-tuning usually takes much less time than pre-training.

For sequence-level classification (e.g., sentence classification or sentence-pair classification), the final hidden state of the special token “[CLS]” is passed to the classifier. For token-level classification, the final hidden state of the token we are interested in is passed to the classifier. In usual, the classifier is a simple neural network with only the input layer and the output layer.

Although BERT has achieved excellent performance, RoBERTa [2] proposes a more robustly optimized approach for BERT pre-training, which leads to a significant improvement in terms of performance in multiple benchmarks. RoBERTa indicates that the performance of BERT model can be improved by: (i) training the model with larger batch size and with more steps, (ii) removing the next sentence prediction task and performing the masked token prediction only, (iii) training the model with longer sequence and (iv) masking tokens dynamically.

BERT-style *pre-train—fine-tune* scheme is ideal for DNA understanding. First, most of the traditional bioinformatics tools develop the understanding of DNA from scratch with task-specific data. As the deep learning models become gradually deeper and wider, their demand for data is getting much more intense. Thus, simply relying on labeled data is very likely to result in poor performance when dataset size is small. In contrast, BERT-style *pre-train—fine-tune* scheme ingeniously utilizes the massive amount of unlabeled data to gain understanding without the need for any human guidance, while such understanding it obtained is easily transferable to various downstream tasks. Therefore, the model can still achieve exceptional performance in data-scarce scenarios. Second, comparing to convolutional neural network (CNN) which only captures local context, Transformer globally capture contextual information from the entire input sequence by taking all the representations from the last layer as input and performing self-attention on them. With the self-attention mechanism, Transformer is not only straightforwardly parallelizable, but also effectively overcomes the gradient vanishing problem that RNN-based architectures usually meet. We believe the reasons above suggests that BERT model has strong potential to lead to many biological breakthroughs if a general understanding of DNA could be formed.

**Pre-training**

Since human genomes are much longer than 512, we used two methods to generate training data. First, we directly split a complete human genome into non-overlapping sub-sequences. Second, we randomly sampled sub-sequences from a complete human genome. The length of each sub-sequence lies in the range of 5 and 510. Specifically, with a 50% probability, we set the length of a sub-sequence as 510. With another 50% probability, we set its length as a random integer between 5 and 510. We regarded each sub-sequence as an independent sequence to train the model. We only perform masked token prediction (i.e. masked language model) in the pre-training step. Unlike natural languages, the unique grammar of the k-mer representation introduces an issue in token masking since a masked k-mer can be trivially made up based on its previous and next k-mers. Taking a 3-mer sequence {CAT, ATG, TGA, GAC, ACT} as an example, “TGA” is the concatenation of “TG” in “ATG” and “A” in “GAC.” If we mask 15 percent of k-mers independently, in most cases, the previous and next k-mers of a masked one are unmasked. This will significantly simplify the pre-training task and prevent the model from learning deep semantic relations among a DNA sequence. Therefore, instead of independently masking each k-mer, we mask contiguous k-length spans of k-mers. For DNABERT we independently feed the last hidden state of each masked token into a classification layer and perform the classification over all the vocabulary. The number of classes here equals to the number of tokens in the vocabulary. At each step, a cross-entropy loss is calculated over all the masked k-mers. We optimize DNABERT with AdamW using the following parameters: $\beta_{1}=0.9, \beta_{2}=0.98, \epsilon=1e-6$ and weight decay as 0.01.

**Fine-tuning**

A majority of DNA applications can be easily formulated as two types: sequence-level task and token-level task. For example, promoter detection can be formulated as a sequence-level 2-class classification task, where class 1 stands for “there is a promoter in the given DNA sequence,” and class 2 stands for the opposite. Moreover, masked token prediction can be formulated as a token-level V-class classification, where V equals the vocabulary size, and each class stands for a unique token in the vocabulary.

DNABERT can solve both the token-level and sequence-level tasks by fine-tuning on the task-specific data. Varying on different tasks, the input sequence can be either a single DNA sequence or the concatenation of multiple sequences. As mentioned above, the output of each layer is a matrix, where each line stands for the representation of its corresponding token. Thus, by feeding the sequence of embeddings to DNABERT, we obtain $L$ matrix, where $L$ is the number of layers. We regard the last matrix (output of the last layer) as the final representation of all the tokens.

For sequence-level tasks, we feed the vector corresponding to the [CLS] token to the output layer. For token-level tasks, we independently feed the vectors corresponding to all the interested tokens to the output layer. Since the DNABERT already captures the contextual and semantic information, the output layer is always as simple as a single layer neural network. Starting from the pre-trained parameters, we exploit labeled data to optimize the DNABERT and the output layer simultaneously. After fine-tuning, the combination of them is capable of solving a specific task. Since the DNABERT only takes sequences shorter than 512, for a longer sequence, we split it into multiple pieces (with a max length of 512), independently feed the pieces to DNABERT, concatenate their final representations together, and feed the concatenated vector to the output layer. We call this model DNABERT-XL, specifically designed to handle longer sequence input.

**Promoter prediction**

We fine-tuned our model DNABERT-Prom using human TATA and non-TATA promoter dataset from the latest version of Eukaryotic Promoter Database (EPDnew), which is a well-annotated non-redundant collection of eukaryotic Pol II promoters that was proven to have high quality [3] (https://epd.epfl.ch/human/human_database.php?db=human). We downloaded 3,065 human TATA and 26,533 non-TATA promoter-containing sequences ranging from -5,000 to +5,000 bp, with +1 being position of transcription start site (TSS). In order to perform benchmarking studies with different existing tools, we trained our binary classification model in two settings. The first setting (hereby referred to as DNABERT-Prom-300) uses 300-bp-long promoter sequences extracted from -249~+50 bp around TSS position as positive class. For the negative set (i.e. non-promoters), simple use of random sequences is not sufficient in ensuring the precision and generalizability since the false discovery rate will be high, as previously discussed in different studies [4, 5]. In order to overcome this issue, we constructed the negative set separately for TATA and non-TATA promoters as follows: for TATA promoters, we randomly picked 3,065 of 300-bp genomic regions not within the -249~+50 bp range but contains the TATA motif. In order to ensure the negative TATA sequence to be as similar to TATA promoters as possible, we made the TATA motif located exactly the same location relative to the actual TATA box (~25 bp upstream of TSS). This way, we forced the model to learn less obvious features and discriminate only through developing understanding of context. For non-TATA promoters, since it does not have the single discriminative feature, we adopted the random substitution approach proposed in [5] with same setting. We found that the dataset constructed this way maintains a good balance between quality and efficiency of data generation, while being more challenging for model to learn. We thus also extended this setting to the core promoter identification by extracting the center 70 bp (-34~+35 bp) sub-sequences, where we trained our model using TATA and non-TATA core promoters altogether while predict separately on TATA and non-TATA datasets. The second setting (DNABERT-Prom-scan) mimics real-world situations, where we scan very long genomic regions with a sliding window and obtain 1001-bp-long sequences for promoter prediction. Naturally, this task is much more difficult than the first setting, given the highly imbalanced (long-tailed) nature of the dataset. We scanned all 10,000-bp-long sequences from EPDnew with a step size of 100 and adopted similar evaluating criteria as in [4]. That is, if the predicted sequence has ≥ 50% (i.e. 500 bp) overlap with -500 to +500 bp region of TSS, it is counted as a TP, otherwise it is counted as a FP. Similarly, failure to make prediction in the area of -500 to +500 bp of TSS will be counted as a FN. Since none of the other methods we found for this task actually provided publicly available scripts for re-training the baselines, we simply used 90% of total data in each setting for training and compare the performances on the remaining 10% test set. Note that this unfairly puts our model at a disadvantage, as observations in our test set may have appeared in other methods’ training data.

**Transcription factor binding sites prediction**

We accessed ENCODE database and obtained the 690 ChIP-seq experiments dataset from UCSC genome browser for fine-tuning of our DNABERT-TF model [6, 7] (<http://hgdownload.cse.ucsc.edu/>goldenPath/hg19/encodeDCC/wgEncodeAwgTfbsUniform/). The ENCODE 690 ChIP-seq dataset covers 161 transcription factor binding profiles in 91 human cell lines and serves as the most popular benchmarking dataset in many TFBS motif prediction studies, such as DeepBind [8], DeepSEA [9], Basset [10], DeepSite [11], DanQ [12] and DESSO [13]. In order to maintain consistency with all the baselines, we extracted the centering 101-bp region around each ChIP-seq peak as positive set. For the negative set, we used similar approach as DESSO, where we pick actual 101-bp long sequences not overlapping with any peaks with same GC content, as we believe the negative sequences created in this manner better resembles the reality [13]. We performed binary classification for the 690 datasets separately using the top 500 even-numbered peaks as test set, following DeepBind [8]. Averaged performances over the 690 datasets were benchmarked with all other tools, which were re-trained following the model specifications in the respective papers, with the best model taken in each case. For analysis of p53, TAp73-alpha and TAp73-beta binding sites, we obtained respective ChIP-seq peaks from Gene Expression Omnibus (GEO) GSE15780 [14] and used those as target regions merged with the p73/p53 binding sites previously predicted by our P53Scan program [15]. The result ~35 bp dimer sequences (binding sites validated by actual ChIP-seq data) were used as input to our model representing positive class. The negative class was built by selecting the top $m$lowest binding site predictions which do not overlap with any ChIP-seq peaks, where $m$ denotes the number of positive sequences for the respective TF. We trained 3 separate models with for the 3 TFs with training and testing set ratio = 9:1.

**Predictions of transcription factor binding sites in mouse genome**

We accessed mouse ENCODE data [16] stored on UCSC genome browser and obtained transcription factor binding sites by ChIP-seq data from Stanford/Yale. Unlike human TFBS data, the mouse TFBS data generated across different labs was not uniformly harmonized so we chose the largest collection available (Stanford/Yale group). We obtained N=78 set of ChIP-seq .narrowPeak files with different antibodies and within different cellular conditions. All settings for data preparation and model training remained the same as those for ENCODE 690 ChIP-seq dataset, except that simple dinucleotide shuffling preserving the frequencies was used for creation of negative set, and that the test set is randomly selected 10% of data instead of top 500 even-numbered peaks.

**Splice donor and acceptor sites prediction**

We followed the same strategy of SpliceFinder [17] in constructing our dataset for splice sites prediction. Specifically, we downloaded human reference genome assembly GRCh38 FASTA file from Ensembl release 99 [18] and extracted 400-bp-long sequences around the donor and acceptor sites of randomly selected exons as positive sequences following the best setting in [17]. The initial dataset consists of equal number (n=10,000) of donor, acceptor and non-splice site sequences, which are non-overlapping intermediate sequences between exons. Nevertheless, this initial setting is found insufficient to detect the non-canonical splice sites without GT or AG dimers. As such, we also reconstructed the dataset adopting an iterative data augmentation approach suggested by [17]. That is, we repeatedly built new models to predict on sliding-window-based scans of previously unseen long genomic sequences. A prediction will only be TP if the splice site is located exactly at the center of the scan. All the false positives were added to negative set and a new model was trained. This process was performed for 100 iterations until the number of false positives in prediction is low. For both initial and reconstructed dataset, we used 90% of data to train our model and tested on the remaining 10% held-out set. In addition, we also constructed a new independent test set with all long genomic sequences not used in our iterative training process, including 114,599 splice site sequences and equal number of randomly picked non-splice site sliding-window scans. We benchmarked with other tools on both the test sets for our initial and reconstructed dataset as well as the independent dataset.

**Evaluation metrics**

For all of the fine-tuning tasks above, the performances were measured in the following metrics (TP = true positive, TN = true negative, FP = false positive, FN = false negative): $Accuracy= \frac{TP+TN}{TP+TN+FP+FN}$; $Precision= \frac{TP}{TP+FP}$; $Recall= \frac{TP}{TP+FN}$; $F1= 2\times\frac{Precision\times Recall}{Precision+Recall}$; $MCC= \frac{TP\times TN-FP\times FN}{\sqrt{(TP+FP)(TP+FN)(TN+FP)(TN+FN)}}$ whenever available. In addition, for the tasks where all selected baselines are able to output continuous probabilities, area under receiver operating characteristics curve (AUROC) and/or area under precision-recall curve (AUPR) were calculated. For three-class classification in splice sites prediction task, average performance metrics (except AUPR) were computed by finding the unweighted mean of performance for each label. The AUROC calculated in this case is the average of all pairwise combinations of classes. ROC and PR curves were plotted separately for splice donor and acceptor prediction in a one-vs-all setting.

**Statistical analysis**

For boxplots in Figure 3a, we applied Wilcoxon one-sided signed-rank test (n=690) for pairwise comparison of the mean and adjusted the p-value using Benjamini-Hochberg procedure. For testing the null hypothesis that all models show equal mean performance (they originate from same distribution), we applied Kruskal-Wallis test, which is a nonparametric version of one-way ANOVA. The statistical significance of differences between AUROCs of models is determined by Delong test, which is a nonparametric method to first compute empirical AUC equivalent to Mann-Whitney statistic by trapezoid rule, and then perform z-test in a paired design [19]. We also applied Mcnemar’s test on the 2×2 contingency table of correct classification vs. misclassification between two classifiers, which is a paired nonparametric test for the null hypothesis that two models have equal performance in terms of proportion of errors on test set.

**Supplementary Figures**

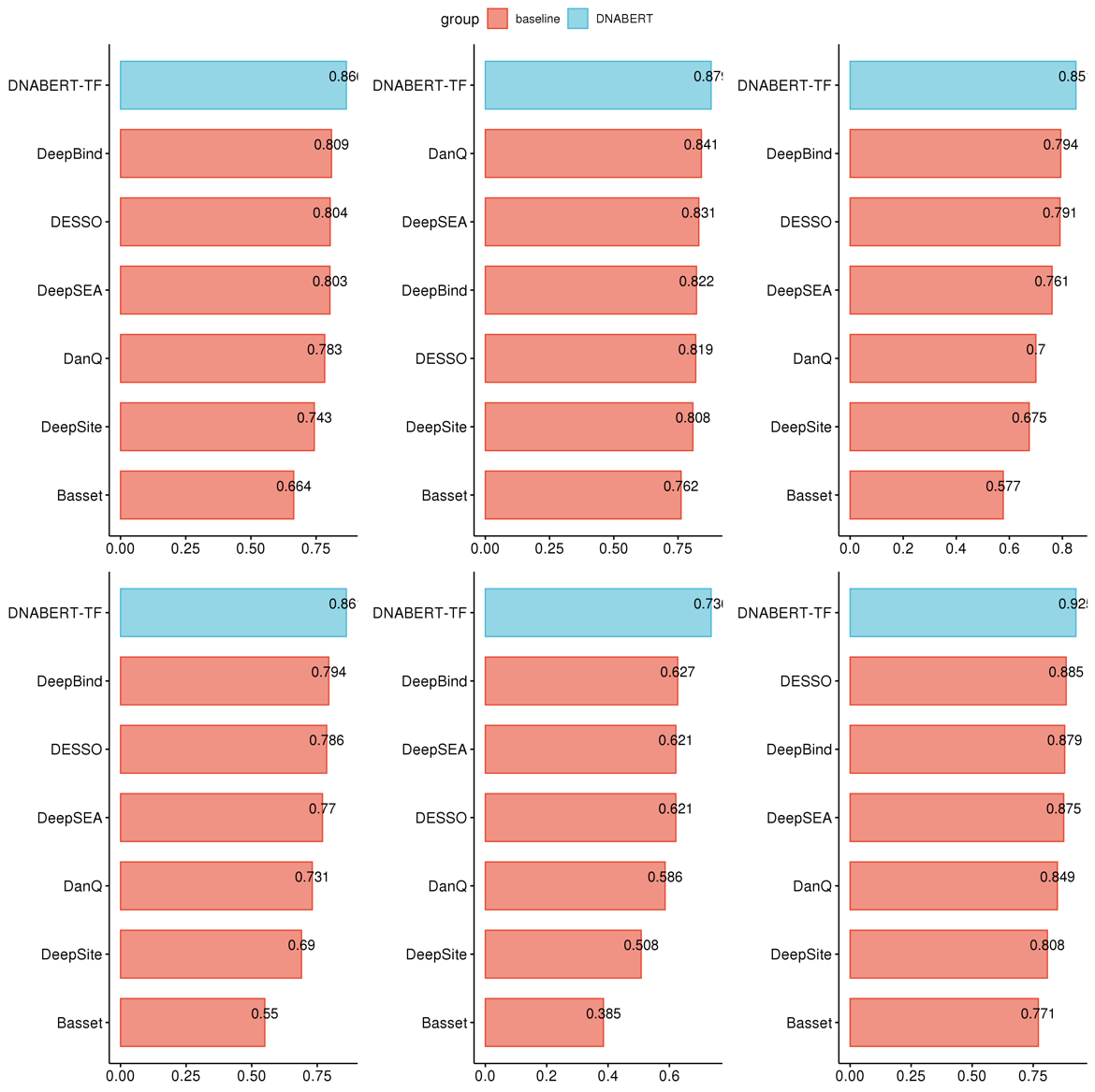

**Supplementary Figure S1.** Performance on ENCODE 690 ChIP-Seq TFBS data (experiments with limited data only). Barplots showing (left to right, top to bottom) accuracy, precision, recall, F1-score, MCC and AUC of DNABERT-TF performance in comparison with other models. ChIP-Seq experiments with limited data is defined as those with less than 10,000 peaks.

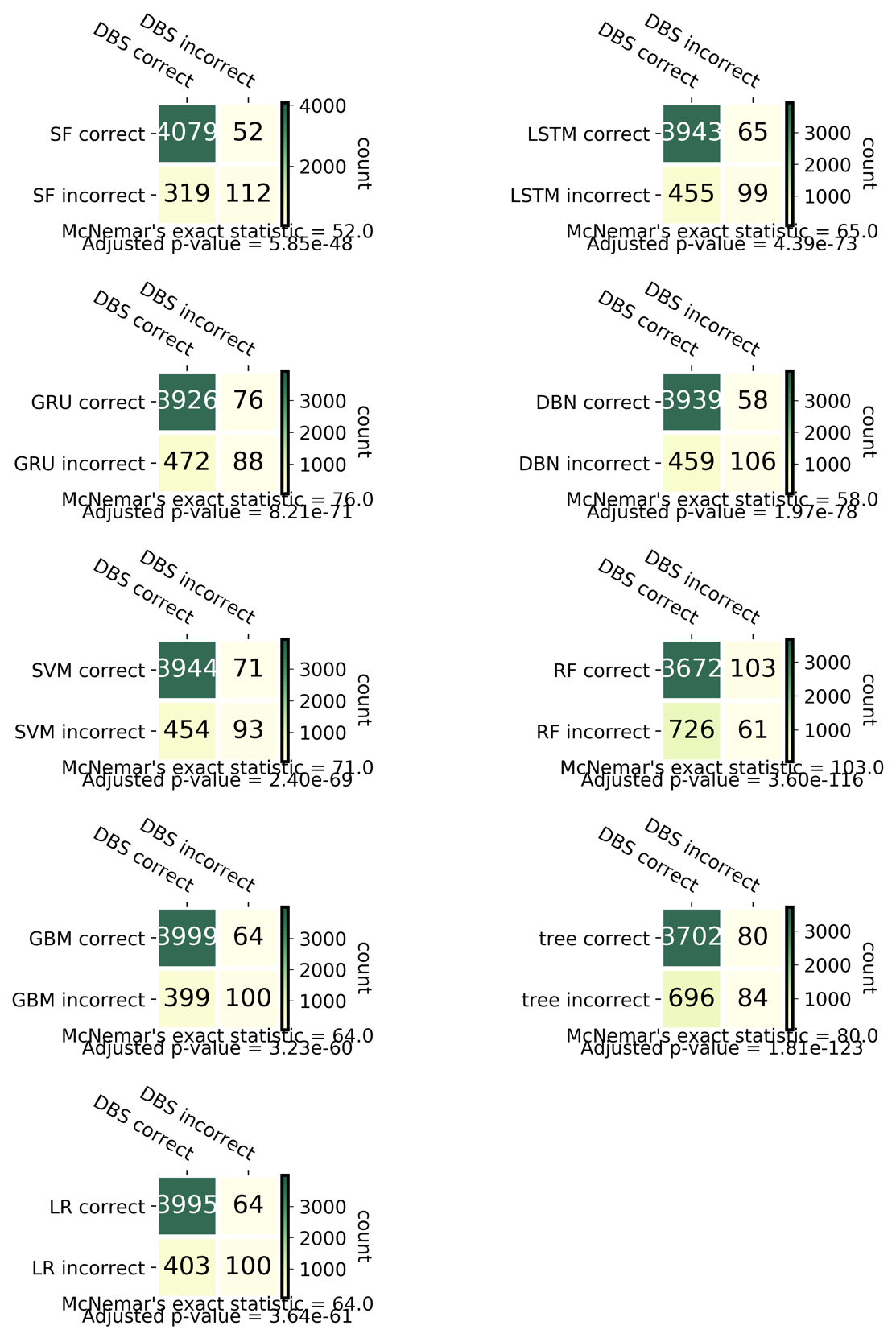

**Supplementary Figure S2.** McNemar’s test between DNABERT-Splice (DBS) and other baseline models on classifying splice donors. SF: SpliceFinder; LSTM: long short-term memory network; GRU: gated recurrent units network; GBM: gradient boosted trees; LR: logistic regression; DBN: deep belief network; RF: random forest; tree: decision tree; SVM_RBF: support vector machine with radial basis function kernel.

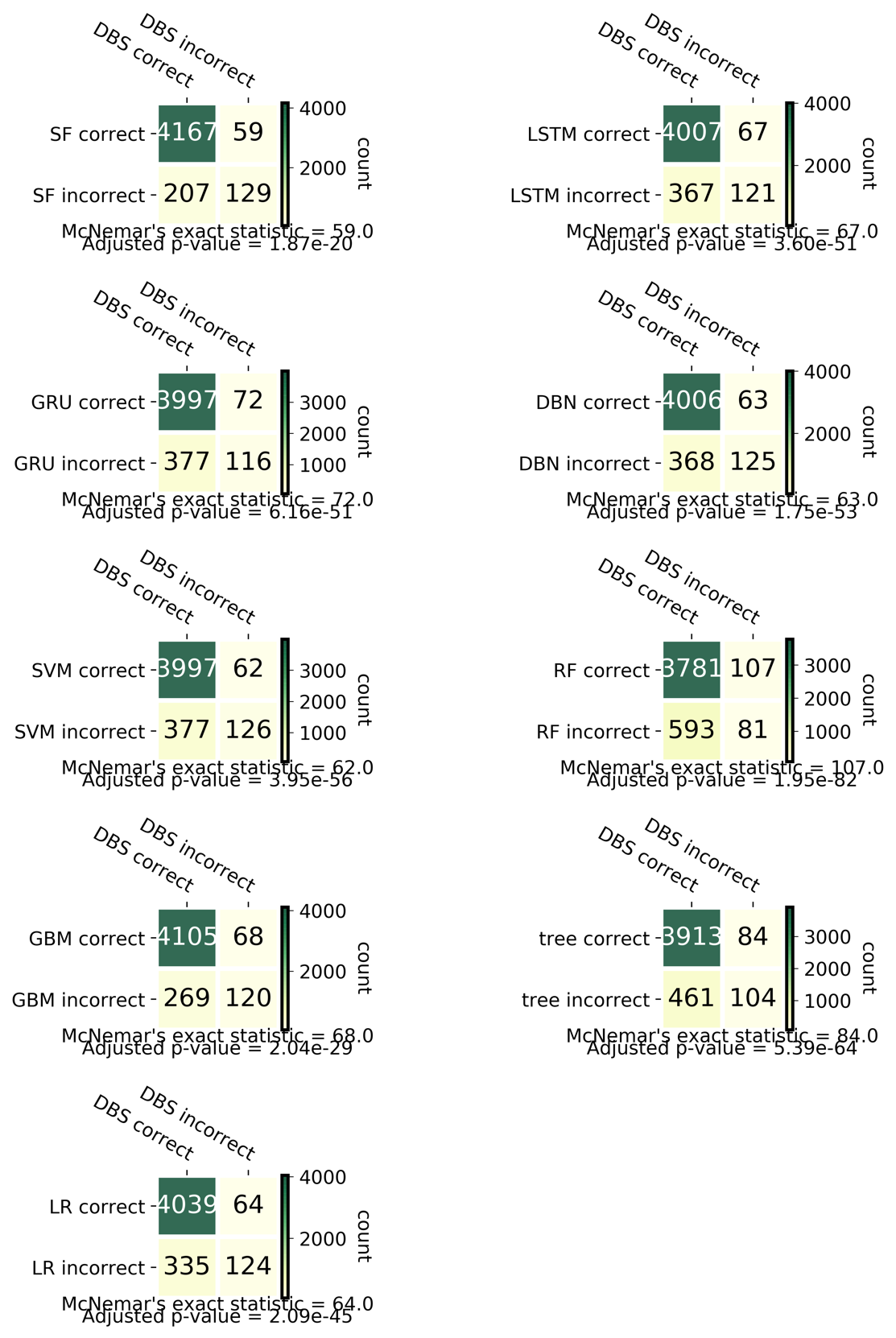

**Supplementary Figure S3.** McNemar’s test between DNABERT-Splice (DBS) and other baseline models on classifying splice acceptors. SF: SpliceFinder; LSTM: long short-term memory network; GRU: gated recurrent units network; GBM: gradient boosted trees; LR: logistic regression; DBN: deep belief network; RF: random forest; tree: decision tree; SVM_RBF: support vector machine with radial basis function kernel.

**Legends for Supplementary Tables**

**Supplementary Table S1.** Mean performance of DNABERT (with kmer = 3, 4, 5, 6) in comparison with six baseline models across six evaluation metrics on ENCODE 690 ChIP-Seq human TFBS data.

**Supplementary Table S2.** Performance of DNABERT (with kmer = 3, 4, 5, 6) on binary classification of p53, TAp73-alpha and TAp73-beta (binding site vs. non-binding site), and DNABERT-6 (kmer = 6) on 3-class classification of TAp73-alpha, TAp73-beta vs. non-binding site across different evaluation metrics.

**Supplementary Table S3.** Performance of DNABERT (with kmer = 3, 4, 5, 6) in comparison with nine baseline models across six evaluation metrics on validation set of initial splice donor and acceptor dataset.

**Supplementary Table S4.** Performance of DNABERT (with kmer = 3, 4, 5, 6) in comparison with nine baseline models across six evaluation metrics on independent splice donor and acceptor test set with model trained on reconstructed dataset.

**Supplementary Table S5.** Hyperparameter settings used for fine-tuning DNABERT models in different tasks.

**Supplementary Table S6.** Complete list of dbSNP Common variants located within high attention regions using input sequences of Prom-300 TATA and noTATA promoter data. Variants with absolute score difference greater than the mean were kept. EPDnew_id: the promoter id used in EPDnew database; coord: the genomic coordinate of 300 bp promoter region; strand: strand of the promoter; gene: gene name (with promoter index); TATA: whether this promoter is a TATA promoter, 1=Yes and 0=No; seq_ori: original sequence with reference allele; seq_mut: altered sequence with alternative allele; SNP: dbSNP rs id; alt_allele: the alternative allele of this variant; pred: prediction with original sequence; pred_mut: prediction with alternative sequence; diff: score difference between pred_mut and pred; log_odds_ratio: log odds ratio between pred and pred_mut; abs_diff: absolute value of diff; is_functional: whether this SNP is documented in ClinVar, GRASP or GWAS Catalog databases, 1=Yes and 0=No.
